# Weight-bearing symmetry changes after asymmetric surface stiffness walking

**DOI:** 10.1101/2025.05.16.654517

**Authors:** Mark Price, Elena G. Schell, Jonaz Moreno Jaramillo, Jenna M. Chiasson, Leah C. Metsker, Meghan E. Huber, Wouter Hoogkamer

## Abstract

Stroke is a leading cause of adult disability in the United States, often presenting as hemiparesis. A priority in hemiparetic gait rehabilitation is the restoration of gait symmetry. While split belt treadmill training has shown promise in correcting spatial gait asymmetry, weight bearing and propulsion asymmetry remain resistant to improvement. As an alternative approach, we tested asymmetric surface stiffness walking to induce signatures of neuromotor adaptation relevant to correcting weight bearing and propulsion asymmetries in hemiparetic stroke. We hypothesized that a bout of asymmetric stiffness walking would elicit aftereffects in the form of asymmetries in weight bearing, propulsion, and plantar flexor activity. Twelve healthy young adults performed a 10-minute bout of asymmetric stiffness walking on an adjustable stiffness treadmill. We measured baseline and post-perturbation ground reaction forces (GRF) and spatio-temporal measures during 5-minute walking bouts on a dual-belt instrumented treadmill. After asymmetric surface stiffness walking, participants walked with increased vertical GRF and plantar flexor muscle excitations during push-off on the perturbed side relative to unperturbed. Participants also decreased their mid-stance vertical GRF and increased their peak braking GRF on the perturbed side relative to unperturbed. Counter to our hypothesis, they did not increase their propulsion GRF on the perturbed side. We conclude that asymmetric stiffness walking elicited a neuromotor adaptation towards a relative increase in push-off in the target limb, albeit primarily vertically aligned in our cohort of healthy young adults, and that gait adaptation to asymmetric stiffness walking should be investigated in individuals with push-off asymmetries.

**New & Noteworthy:** Weight-bearing asymmetry in individuals with hemiparetic stroke is resistant to treatments that produce improvements in other gait function measures (e.g., spatio-temporal symmetry). We investigated a novel perturbed ground stiffness intervention applied by an adjustable surface stiffness treadmill and found significant aftereffects in vertical and braking ground reaction force peaks in healthy young adults, as well as increases in perturbed-side plantar flexor activity during push-off, indicating a strong potential for correcting persistent deficiencies in post-stroke gait.

## Introduction

Stroke is the leading cause of adult disability in the United States (1) and leaves 80% of its survivors with gait impairments (2), many of which result from gait asymmetries due to hemiparesis (3–5). Hemiparetic stroke, or stroke resulting in motor deficits on one side of the body, typically causes weakness in the lower limb muscles on the affected side. Weakened dorsiflexors are associated with increased fall risk during walking due to insufficient dorsiflexion during swing (e.g., drop-foot) (6). Weakened plantar flexors are associated with reduced propulsion, which is theorized to be a major contributor to several common gait deficits resulting from stroke (7, 8), such as slower walking speed (9, 10), asymmetric step lengths (11), asymmetric step time (12), and asymmetric weight bearing (13). Of particular concern, asymmetric weight bearing during walking can eventually increase risk of osteoarthritis (14) and osteoporosis (15).

Split-belt treadmill training is a recent form of physical therapy shown to reduce gait asymmetries in hemiparetic individuals post-stroke (16). In split-belt treadmill training, a participant walks on treadmill with two belts, one for each leg. The belt speeds are different, set to exaggerate the asymmetry in step length, which drives neuromotor adaptation towards less asymmetric step lengths after exposure (17). While repetitive bouts of such short-term adaptation lead to longer-term improvements in gait asymmetry (16), a major shortcoming of current split-belt treadmill training protocols is that they do not reduce temporal asymmetries that result from asymmetries in weight-bearing and propulsion forces (18, 19). Alternatively, real-time audiovisual biofeedback displaying real-time propulsive ground reaction forces (GRFs) or trailing limb angles can improve propulsion (20, 21), but the benefits of such explicit learning interventions might not result in long-lasting benefits as observed for implicit learning interventions (22).

Asymmetric surface stiffness treadmill walking has been shown to change gait patterns in healthy individuals using implicit locomotor adaptation principles similar to those observed in split-belt treadmill walking (23). In this paradigm, rather than exposing participants to asymmetrical belt speeds, they are instead exposed to asymmetrical ground stiffness, during which the platform under one foot deflects downward in response to walking loads (23). Because the perturbation in this case is responsive to the limb loading dynamics of gait, it is theorized that it could elicit adaptations to weight-bearing and propulsion force symmetry. Encouragingly, vertical GRF during push-off seemed to increase by 17% in healthy young adults after asymmetric stiffness walking (23) although reported vertical GRF values were lower than typically observed (24). Despite these preliminary results, it is unclear if weight-bearing or propulsion symmetry are affected by asymmetric stiffness walking, as vertical GRFs were measured only for one side, and propulsion GRFs were not measured at all. Furthermore, while muscle activity was measured, it was calculated only for the swing phase of walking, despite changes in plantar flexor activity during push-off being of critical importance for determining the promise of this technique for addressing hemiparetic deficits (8).

There is thus a critical need to quantify neuromotor adaptation to asymmetric stiffness walking in vertical and horizontal GRFs, muscle activity, and spatiotemporal gait parameters. This study aims to 1: quantify the bilateral effects of neuromotor adaptation to an asymmetric stiffness walking intervention in measures relevant to common gait manifestations in post-stroke gait, and 2: understand mechanisms that underlie how asymmetric stiffness walking produces aftereffects in weight bearing and muscle activity in healthy individuals. Extrapolating from the preliminary findings in (23), we hypothesize that after a single bout of asymmetric surface stiffness walking, vertical (vGRF) and propulsive ground reaction forces (pGRF), and stance time will increase on the low stiffness side relative to the unperturbed side. Corresponding with these changes, we hypothesize that gastrocnemius and soleus muscle excitation will increase during push-off on the low stiffness side relative to the unperturbed side. We also hypothesize that tibialis anterior activity will increase during the swing phase on the unperturbed side due to the increased relative height of the rigid treadmill during stance on the low stiffness side. In addition to testing these hypotheses, we performed a holistic analysis of GRF and EMG measures.

## Methods

### Participants

Twelve healthy young individuals participated in this study (21 ± 1 years, 69.3 ± 8.1 kg, 164 ± 7 cm, 11F/1M). All participants were free of musculoskeletal or neurological injury, and disease or surgery that would affect gait or balance behavior. They were capable of walking at a comfortable pace for 30 consecutive minutes and had no prior experience walking on an asymmetric surface stiffness treadmill. Participants gave their written informed consent in accordance with the ethical guidelines by the UMass Amherst Institutional Review Board (Protocol 5248).

### AdjuSST

The AdjuSST (Adjustable Surface Stiffness Treadmill) is our custom-built dual belt treadmill with one belt operating around a rigid platform, and the other belt around a platform resting atop a leaf spring mechanism allowing for vertical deflection under weight (25). For this study we operated the adjustable side in two settings: rigid (300 kN/m) and compliant (15 kN/m). In the compliant setting, under a static load of 750 N (∼75 kg) the belt displaces 50 mm downwards. Mechanical stops with shock absorbing foam prevent movement beyond 100 mm. We set the perturbed side to right or left by specifying the walking direction and running the belts in forward or reverse accordingly. The first eight participants experienced a left-side perturbation, and the final four experienced a right-side perturbation, for logistical reasons.

### Protocol

Participants performed two 5-minute walking trials on a force-instrumented dual-belt treadmill (Bertec, Columbus, OH, USA), one before and one after walking a 12-minute walking trial on the AdjuSST, including 10 minutes of walking with one leg at the rigid belt, and one leg at the compliant belt at a stiffness of 15 kN/m (Figure 1). The 5-minute walking trials allowed us to compare baseline walking kinetics and muscle activations to potential after-effects in walking kinetics and muscle activations from the motor adaptation to asymmetric surface stiffness walking.

**Figure 1.**
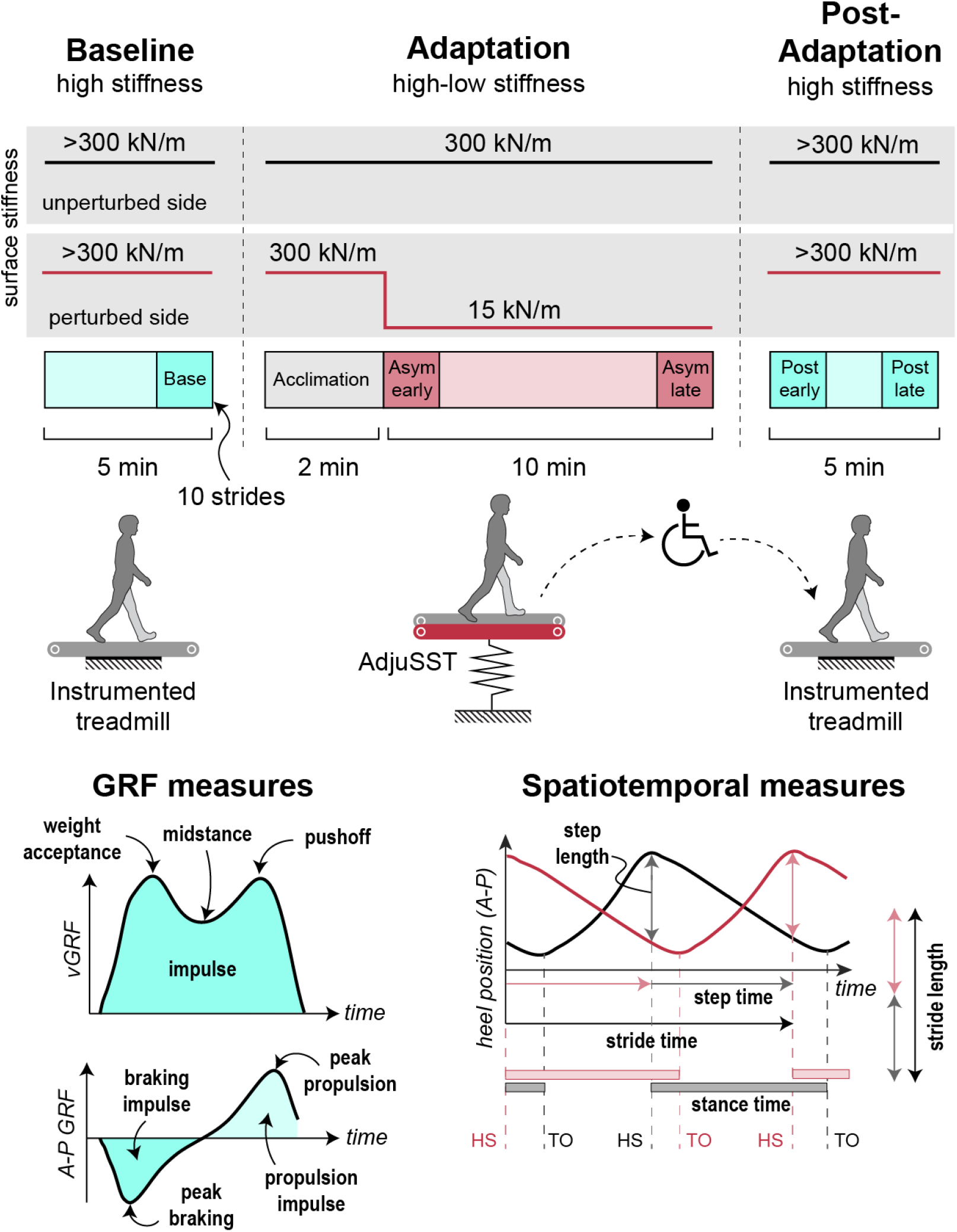
Experimental setup and paradigm. A. Participants experienced asymmetric surface stiffness walking on a custom adjustable stiffness treadmill (AdjuSST; (25)). B. Experimental paradigm. Participants walked in 3 total bouts – a 5-minute baseline bout on a rigid instrumented treadmill, a 12-minute bout on the AdjuSST in which the surface stiffness was abruptly lowered on one side for the latter 10 minutes, and a 5-minute post-perturbation bout on the rigid instrumented treadmill. Participants were transferred to the instrumented treadmill by wheelchair to prevent washout of adaptation effects in transit.

Experimental set up before the first walking trial consisted of fitting the participant with proper sizes of shoes (Speed Sutamina, Puma, Herzogenaurach, Germany) and force measuring insoles (loadsol, Novel, Munich, Germany). We placed ten lower limb surface electromyography (EMG) sensors (Delsys Trigno, Natick, MA, USA)) bilaterally on five lower limb muscles: vastus lateralis (VL), tibialis anterior (TA), gastrocnemius lateral head (GAS), soleus (SOL), and biceps femoris (BF). Following, we placed four markers on the participants’ shoes, one on each heel, and one on each fifth metatarsal head, used for gait event detection during AdjuSST walking. Participants first performed a 5-minute walking trial on the force-instrumented treadmill (“Baseline”). For all conditions, the participants walked at 1.25 m/s. We collected ground reaction force (GRF) data at 1000 Hz, and EMG data at 1926 Hz using QTM (Qualisys, Göteborg, Sweden) and insole force data at 100 Hz for 60 seconds during the final minute of the 5-minute trial.

After the Baseline trial, we led participants to the AdjuSST, located in a separate lab space, ∼60m from the force-instrumented treadmill. Participants then completed a 12-minute walking trial on the AdjuSST. For the first two minutes of this trial, both belts were set at high stiffness, similar to ground stiffness. After 2 minutes, one belt’s stiffness condition was lowered to 15 kN/m within a single step. Participants walked for 10 minutes with one belt set at high stiffness and the other set at 15 kN/m. We measured EMG data at 1926 Hz and foot marker location at 100 Hz with QTM, and insole force data at 100 Hz for 60 seconds during the first and final minute of these 10 minutes. Upon completion of the 10-minute intervention condition we stopped the treadmill and asked participants to not take any additional steps, in efforts to decrease preemptive washout of aftereffects. We then transferred participants into a wheelchair and transported them back to the force-instrumented treadmill.

Participants then performed a 5-minute walking trial on the force-instrumented treadmill (“Post”), which allowed us to quantify aftereffects in gait symmetry. We collected ground reaction force (GRF) data at 1000 Hz, and EMG data at 1926 Hz using QTM and insole force data at 100 Hz for 60 seconds during the first and final minute of the 5-minute trial.

### Data analysis

We smoothed ground reaction force vectors and centers of pressure obtained from the rigid instrumented treadmill with a fourth-order Butterworth low-pass filter with a cut-off frequency of 30 Hz. We identified heel strike and toe-off events for each limb as the moment in which the force vector magnitude for the corresponding force plate exceeded (heel strike) or fell beneath (toe-off) a threshold of 100 N. Gait events were manually verified for all strides, and strides in which the stance foot interacted with both force plates simultaneously were discarded. Motion capture marker trajectories were smoothed with a fourth-order Butterworth low-pass filter with a cutoff frequency of 6 Hz.

Vertical ground reaction forces from the force measuring insoles were initially processed in the same manner as the force plate data from the rigid treadmill. We additionally calibrated the insole measurements against the force plate measurements recorded during the baseline walking trial. We compared the vGRF trajectory averaged across 20 corresponding strides between the force plates and the insoles, then applied a gain and offset to the forefoot and hindfoot insole signals to match the peak values of the resultant vGRF with the force plate vGRF peaks, as well as to zero the insole reading during swing (Supplementary Figure 1S). The gain and offset calculated from this sample were applied to all corresponding insole measurements for the duration of the experiment, per participant.

We calculated muscle excitation profiles from the EMG signals by applying a fourth-order Butterworth band-pass filter (30 – 300 Hz) to eliminate noise in the signal. We then DC bias corrected, rectified, and applied a fourth-order Butterworth low-pass filter (5 Hz) to create the linear envelope for each muscle excitation signal (26, 27). We calculated the peak value of each linear envelope signal per stride, averaged the peaks for the final ten strides of the baseline trial, and used the resulting value to scale the filtered signal for the corresponding sensor for all conditions. Because of sensor malfunctions, 2 of the 12 participants could not be included for any EMG analyses, and 1 additional participant was not included in each of the SOL and the VL analyses. We performed all experimental data processing with custom scripts using MATLAB (Mathworks, Natick, MA, USA).

### Dependent Measures

To quantify weight bearing, we calculated the first (weight acceptance) and second (pushoff) vGRF peak, the minimum vGRF during midstance, and the time-integral of the vGRF profile during stance (vGRF impulse) per side, per stride. To quantify propulsion, we calculated the peak pGRF and pGRF impulse per side, per stride. We also calculated the peak and impulse of the braking ground reaction force (bGRF) per side for all strides.

To quantify spatiotemporal parameters of gait, we calculated stance time per side as the elapsed time between heel strike and subsequent toe off events. We similarly calculated step time per side as the elapsed time between heel strike of the contralateral limb and subsequent heel strike of the ipsilateral limb. We calculated step length as the anterior-posterior distance between heel markers at the moment of heel strike of the corresponding side. Stride time and stride length are defined as the sum of the step time or step length of both sides for a given stride.

To quantify changes in muscle activity corresponding with our hypothesized changes in weight bearing and propulsion, we calculated peak muscle excitations of the gastrocnemius and soleus during push-off (between 30 and 70% of the gait cycle) and the vastus lateralis during weight acceptance (0 to 20%). To further investigate changes relevant to post-stroke gait (i.e., drop foot, knee hyperextension), we calculated peak excitation of the first burst of tibialis anterior activity during swing (60 to 80%) (28) and peak excitation of the biceps femoris during terminal swing (80 to 100%).

For all unilateral measures, we calculated an asymmetry ratio to indicate the relative magnitude of a measure on the perturbed vs. unperturbed side. This ratio is defined as:

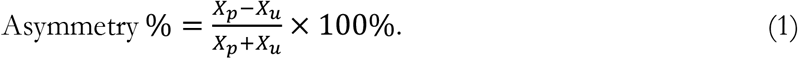

where *X*_*p*_ is the measure calculated for the side that was perturbed with low surface stiffness, and *X*_*u*_ is the measure for the side that remained unperturbed. Unless otherwise specified, all statistical analyses were performed on the asymmetry ratio of the corresponding measure.

### Statistical analyses

We averaged all measures per participant in 10-stride windows to represent three experimental conditions: the final 10 strides of the baseline walking trial (“Baseline”), the initial 10 strides on the instrumented treadmill after exposure to asymmetric surface stiffness (“Post-early”), and the final 10 strides after 5 minutes of post-perturbation walking (“Post-late”). We defined two additional conditions for measures calculated from the insoles and EMG sensors as those we able to be recorded during the perturbation: the first (“Asym-early”) and final (“Asym-late”) 10 strides of asymmetric stiffness walking.

To determine whether asymmetric stiffness walking significantly influenced a given measure, we used a one-way repeated measures analysis of variance (ANOVA). We tested for sphericity (i.e., the assumption of equal variances of the differences between all combinations of conditions) with Mauchly’s test; if the assumption of sphericity was violated, we applied the Greenhouse-Geisser correction factor to the degrees of freedom of the ANOVA. For measures with a significant effect of walking period condition, we conducted a post-hoc comparison using a two-tailed paired t-test between the Baseline and Post-early conditions to test our hypothesis that asymmetric walking would result in an asymmetry relative to baseline after the perturbation (see the introduction for predictions for specific measures). We performed all statistical analyses using MATLAB (Statistics and Machine Learning Toolbox), and the significance level was set at α = 0.05 for all tests.

## Results

### Weight bearing

We found a significant effect of walking period condition in push-off (p<.001) and mid-stance (p<.001) vGRF peaks, but not during weight acceptance (p=.055) or vGRF impulse (p>.1) (Table 1; Figure 2). Relative to Baseline, participants had larger push-off vGRF peaks (+2.8% asymmetry, p<.001) on the perturbed side than the unperturbed side during the Post-early window (Figure 3). Participants had lower mid-stance vGRF on the perturbed than the unperturbed side (-2.2% asymmetry, p<.001) in Post-early relative to Baseline.

**Table 1.**
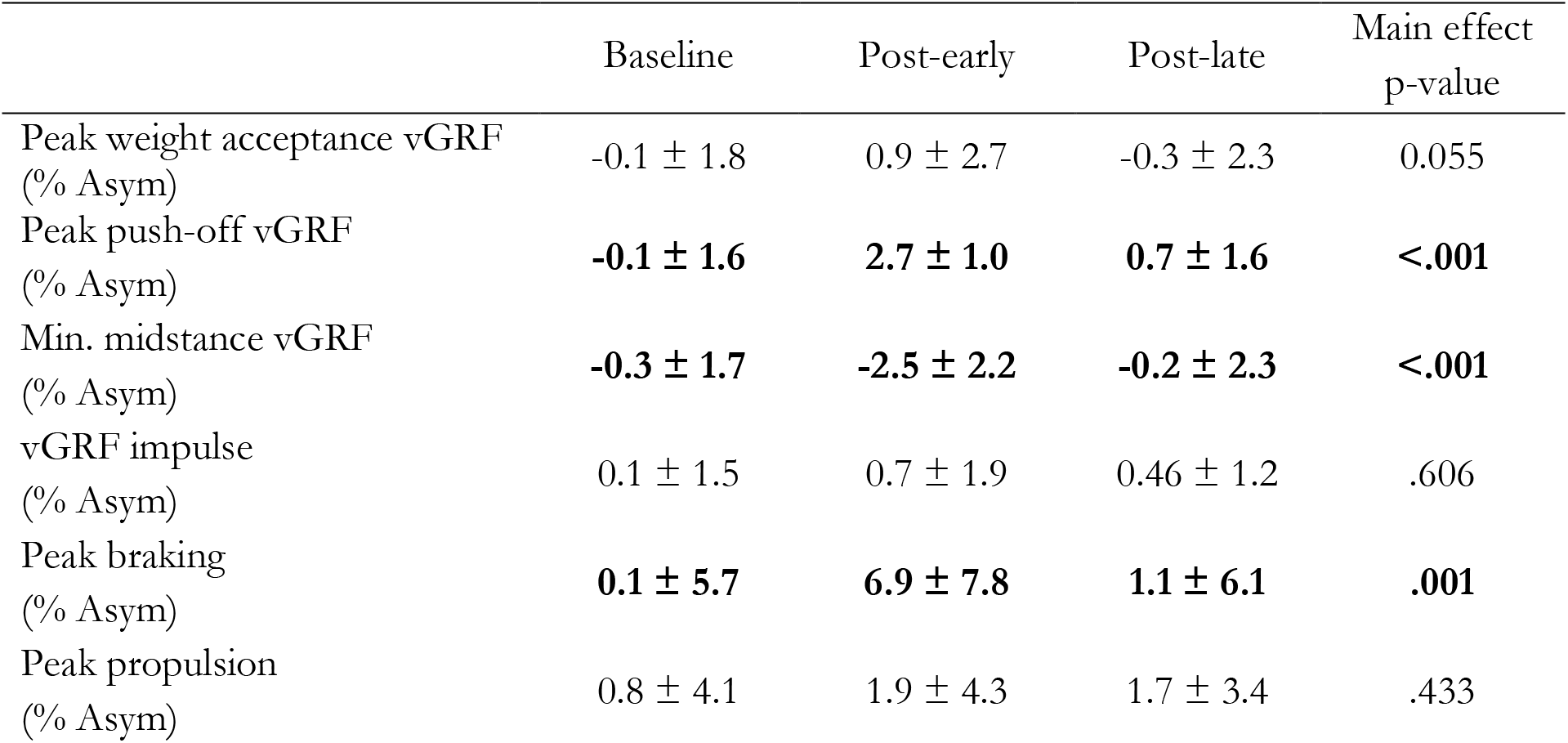
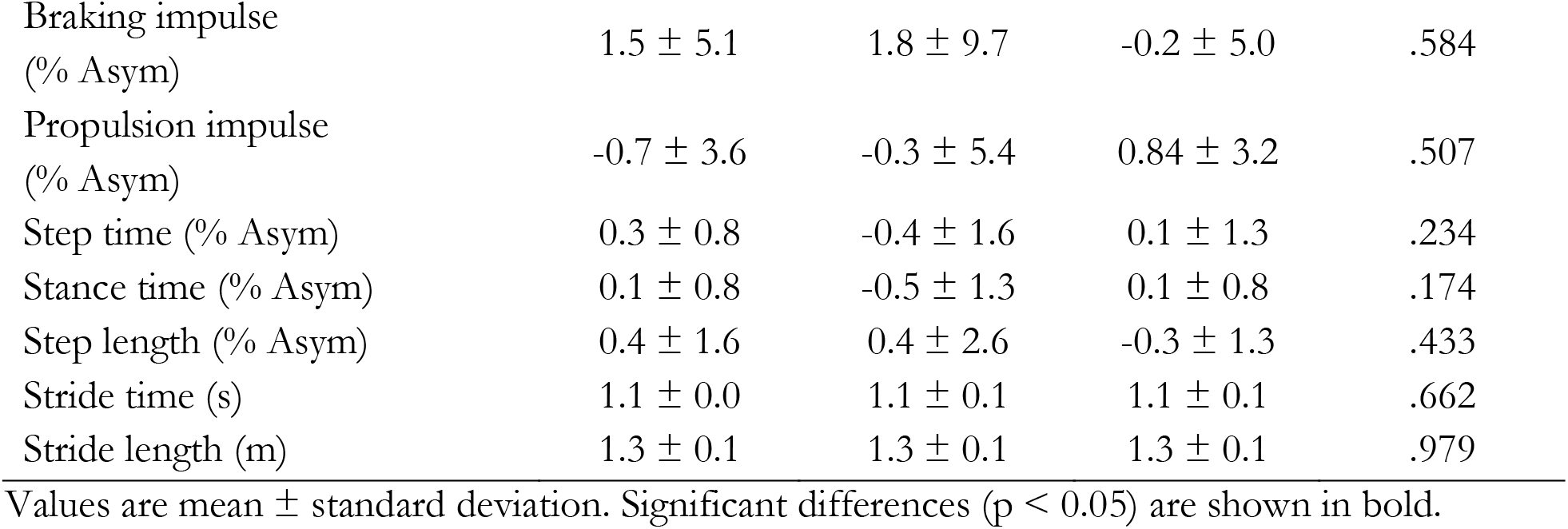
Ground reaction force and spatiotemporal outcome measures.

|  | Baseline | Post-early | Post-late | Main effect<br>p-value |
| --- | --- | --- | --- | --- |
| Peak weight acceptance vGRF<br>(% Asym) | -0.1 ± 1.8 | 0.9 ± 2.7 | -0.3 ± 2.3 | 0.055 |
| Peak push-off vGRF<br>(% Asym) | **-0.1 ± 1.6** | **2.7 ± 1.0** | **0.7 ± 1.6** | **<.001** |
| Min. midstance vGRF<br>(% Asym) | **-0.3 ± 1.7** | **-2.5 ± 2.2** | **-0.2 ± 2.3** | **<.001** |
| vGRF impulse<br>(% Asym) | 0.1 ± 1.5 | 0.7 ± 1.9 | 0.46 ± 1.2 | .606 |
| Peak braking<br>(% Asym) | **0.1 ± 5.7** | **6.9 ± 7.8** | **1.1 ± 6.1** | **.001** |
| Peak propulsion<br>(% Asym) | 0.8 ± 4.1 | 1.9 ± 4.3 | 1.7 ± 3.4 | .433 |
| Braking impulse<br>(% Asym) | 1.5 ± 5.1 | 1.8 ± 9.7 | -0.2 ± 5.0 | .584 |
| Propulsion impulse<br>(% Asym) | -0.7 ± 3.6 | -0.3 ± 5.4 | 0.84 ± 3.2 | .507 |
| Step time (% Asym) | 0.3 ± 0.8 | -0.4 ± 1.6 | 0.1 ± 1.3 | .234 |
| Stance time (% Asym) | 0.1 ± 0.8 | -0.5 ± 1.3 | 0.1 ± 0.8 | .174 |
| Step length (% Asym) | 0.4 ± 1.6 | 0.4 ± 2.6 | -0.3 ± 1.3 | .433 |
| Stride time (s) | 1.1 ± 0.0 | 1.1 ± 0.1 | 1.1 ± 0.1 | .662 |
| Stride length (m) | 1.3 ± 0.1 | 1.3 ± 0.1 | 1.3 ± 0.1 | .979 |

Values are mean ± standard deviation. Significant differences ( $p < 0.05$ ) are shown in bold.

**Figure 2.**
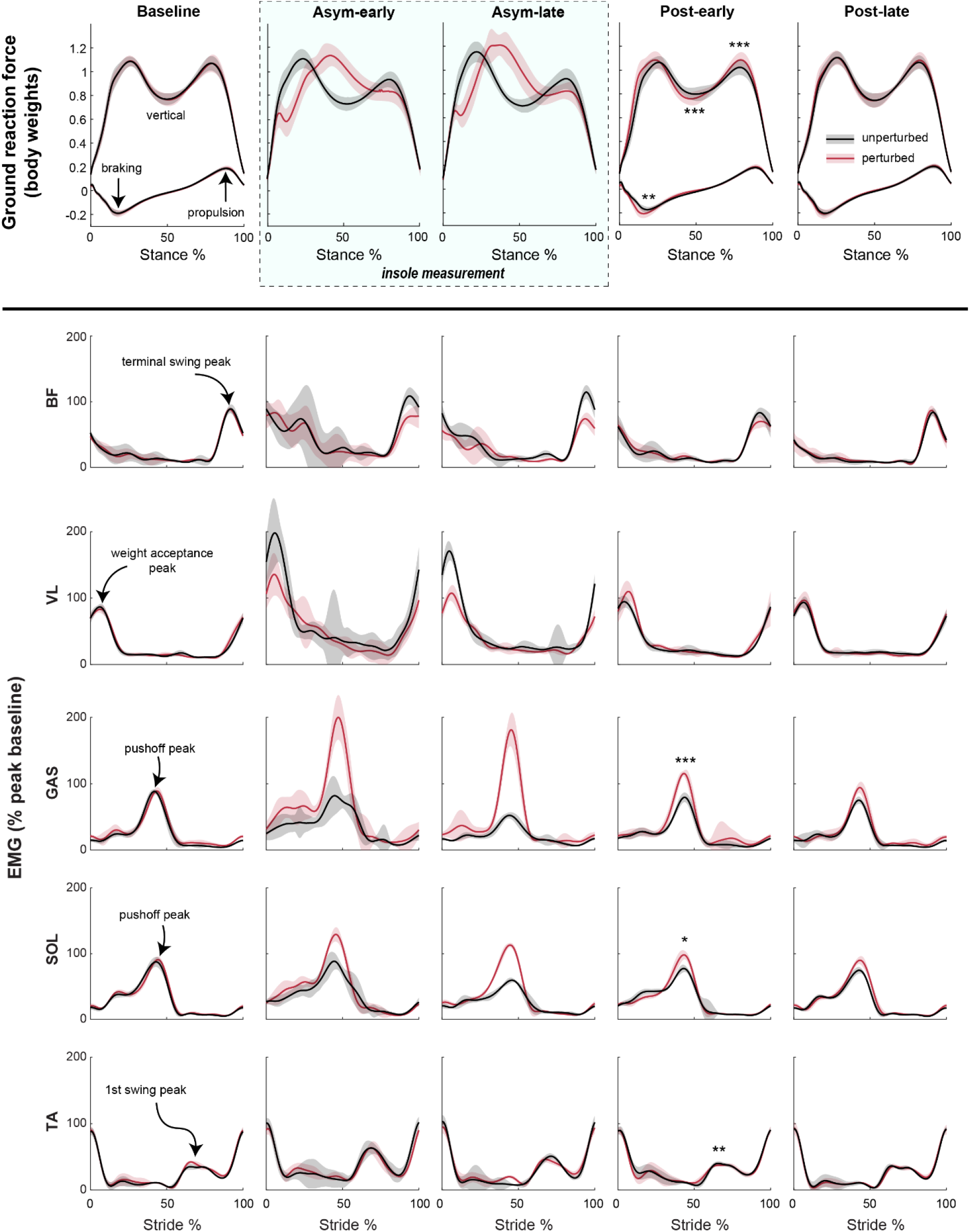
Electromyography (EMG) and ground reaction force (GRF) traces during Baseline, Asym-early, Asym-late, Post-early, and Post-late conditions. Selected measures are illustrated and labeled in the “Baseline” plots. Asterisks reflect significant difference to baseline via paired t-test: ^*^ = p<.05, ^**^ = p<.01, ^***^ = p<.001.

**Figure 3.**
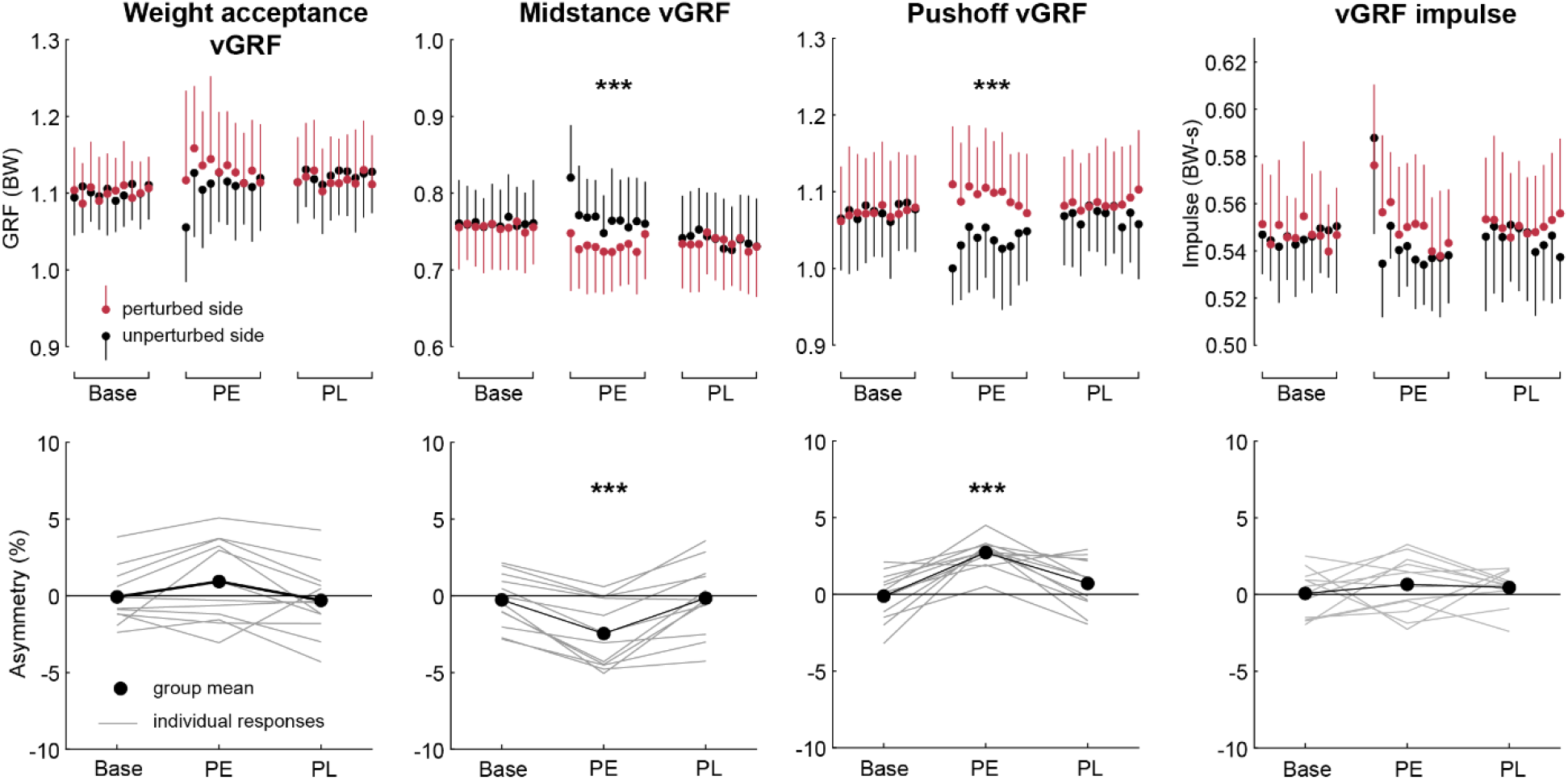
Group means of vertical GRF measures per condition (Base = Baseline, PE = Post-early, PL = Post-late) for (top) the perturbed and unperturbed limbs per stride and (bottom) asymmetry ratio. Error bars reflect one standard deviation. Asterisks reflect significant difference to baseline via paired t-test: ^*^ = p<.05, ^**^ = p<.01, ^***^ = p<.001.

### Propulsion and braking

We found a significant effect of walking period condition in peak braking force (p=.001), but not peak propulsion, braking impulse, or propulsion impulse (Table 1; Figure 2). Relative to Baseline, participants had larger peak braking (+6.8% asymmetry, p=.006) on the perturbed than the unperturbed side during the Post-early window (Figure 4).

**Figure 4.**
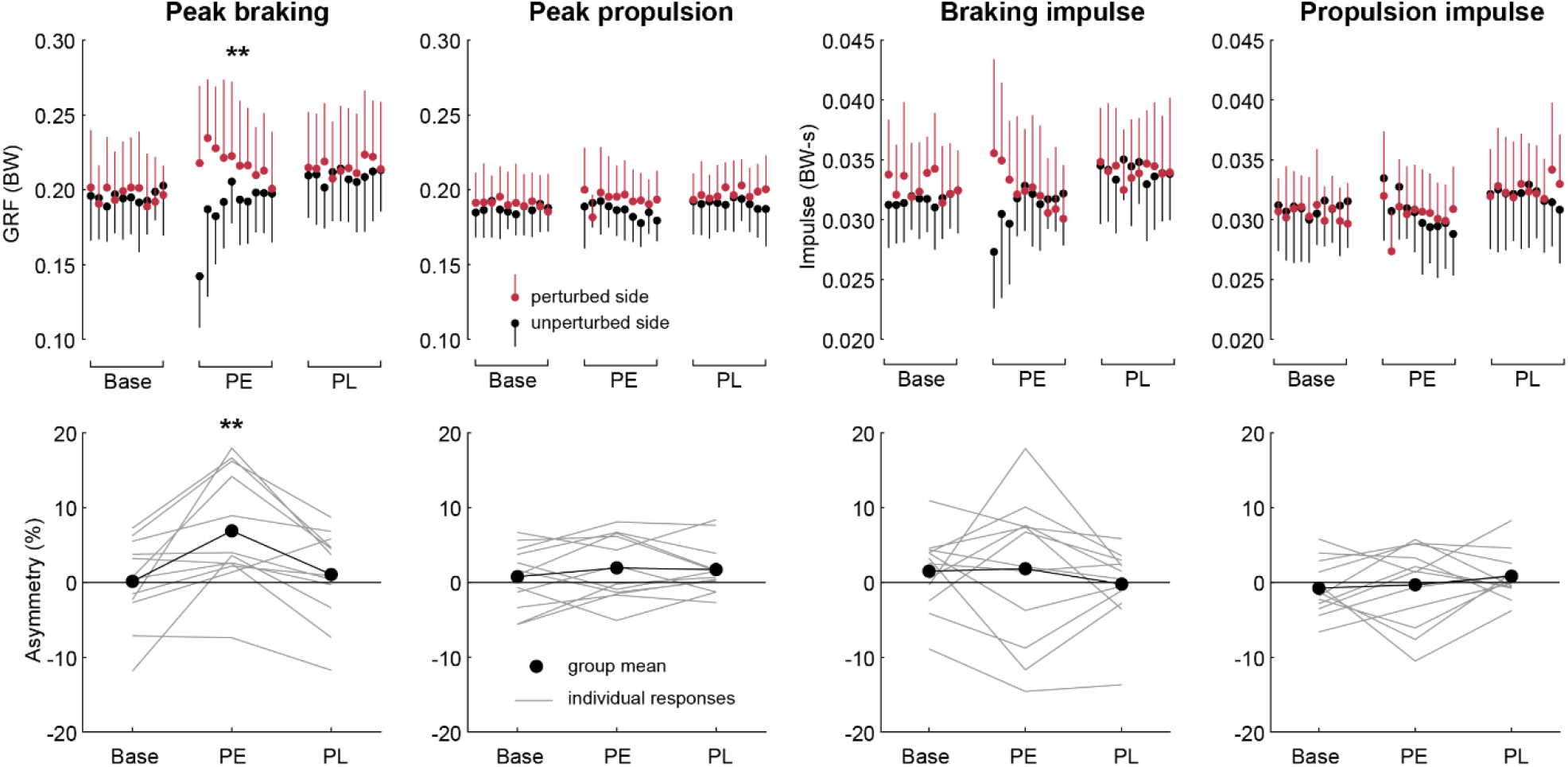
Group means of braking GRF and propulsive GRF measures per condition (Base = Baseline, PE = Post-early, PL = Post-late) for (top) the perturbed and unperturbed limbs per stride and (bottom) asymmetry ratio. Error bars reflect one standard deviation. Asterisks reflect significant difference to baseline via paired t-test: ^*^ = p<.05, ^**^ = p<.01, ^***^ = p<.001.

### Spatiotemporal

We did not find a significant effect of walking period condition in step time symmetry, stance time (symmetry, or overall stride time (p>.1 for all cases) (Table 1). Likewise, we did not find a significant effect of walking period condition in step length or stride length (p > .1) (Supplementary Figure 2S).

### Muscle excitations

We found a significant effect of walking period condition on asymmetry in all five EMG measures (Table 2; Figure 2). Of these, GAS (+20.7% asymmetry, p<.001), SOL (+9.5% asymmetry, p=.016), and TA (-8.8% asymmetry, p = .002) showed asymmetry aftereffects relative to baseline. Both plantar flexors showed relatively higher muscle activity on the perturbed side during the post-late condition, though the TA effect was to relatively increase activity on the unperturbed side, in part canceling out a baseline asymmetry of +5.4% favoring the perturbed side (Figure 5).

**Table 2.** Muscle excitation outcome measures.

|  | Baseline | Asym-early | Asym-late | Post-early | Post-late | Main effect<br>p-value |
| --- | --- | --- | --- | --- | --- | --- |
| BF (% Asym) | **0.4 ± 1.0** | **-12.1 ± 13.3** | **-19.9 ± 21.0** | **-4.8 ± 13.4** | **1.8 ± 17.0** | **.003** |
| VL (% Asym) | **-0.1 ± 2.5** | **-17.2 ± 17.2** | **-20.3 ± 14.4** | **5.3 ± 13.1** | **2.5 ± 9.8** | **< .001** |
| GAS (% Asym) | **-0.8 ± 2.4** | **34.2 ± 18.5** | **47.7 ± 10.2** | **19.9 ± 8.3** | **9.7 ± 8.5** | **< .001** |
| SOL (% Asym) | **0.2 ± 0.9** | **13.9 ± 15.4** | **27.2 ± 12.4** | **9.7 ± 8.8** | **8.9 ± 8.0** | **< .001** |
| TA (% Asym) | **5.3 ± 8.2** | **-4.1 ± 11.8** | **-6.2 ± 13.8** | **-3.5 ± 10.9** | **-1.3 ± 9.0** | **.014** |

Values are mean ± standard deviation. Significant differences ( $p < 0.05$ ) are shown in bold.

**Figure 5.**
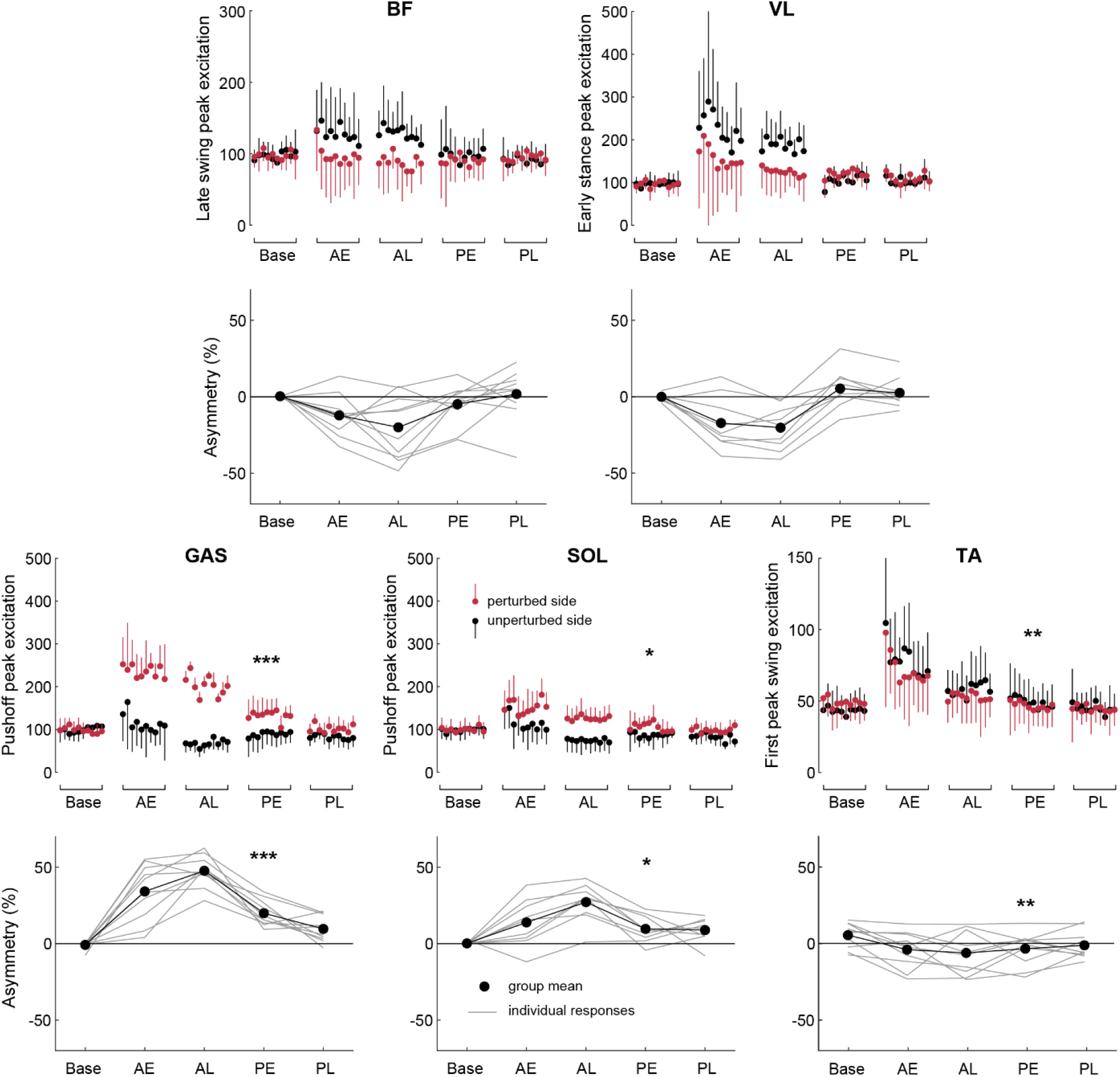
Group means and standard deviations of EMG measures per condition (Base = Baseline, AE = Asym-early, AL = Asym-late, PE = Post-early, PL = Post-late) (top) per stride and (bottom) asymmetry ratio. Error bars reflect one standard deviation. Asterisks reflect significant difference to baseline via paired t-test: ^*^ = p<.05, ^**^ = p<.01, ^***^ = p<.001.

## Discussion

Overall, participants presented significant weight-bearing aftereffects after exposure to asymmetric stiffness walking. In support of our hypothesis, peak push-off vGRF shifted toward the perturbed side limb along with associated muscle excitations in the plantar flexors. Counter to our hypothesis, we did not observe a significant shift in propulsion force, vGRF impulse, or stance time symmetry.

### Weight-bearing adaptation and aftereffects

Participants changed their weight bearing patterns in response to the intervention. During adaptation, the vGRF profile was distorted, losing its “double hump” shape almost completely, with the first peak increasing in magnitude and the second peak nearly disappearing (Figure 2). Interestingly, post-perturbation, the weight acceptance peak was not significantly affected while the push-off peak on the perturbed side substantially increased relative to baseline. This may indicate that the qualitatively large increase in weight acceptance vGRF during the perturbation is mainly due to the passive mechanical dynamics of asymmetric stiffness walking and did not require a major adaptation to the neuromotor control of walking. Visual analysis of per-stride weight acceptance vGRF reveals an initial asymmetry post-perturbation (Figure 3) but appears to wash out within fewer than 10 strides. This rapid washout suggests that re-acclimation to a “normal” walking context occurs on a short timescale, consistent with fast adaptation rather than longer-term learning. In contrast, the reduction or disappearance of the push-off peak during the perturbation appears to have elicited compensations toward increased vertical push-off that we observed in the post-early window. In support of this explanation, we observed post-perturbation increases in activity for muscles that contribute to push-off (GAS and SOL) on the perturbed side, but did not observe aftereffects in muscles that play a greater role in body support (BF and VL). During the late-asymmetry condition, activity appeared to increase on the unperturbed side while remaining relatively unchanged on the perturbed side for both BF and VL. In contrast, GAS and SOL both appeared to increase on the perturbed side during the perturbation, and that increased activity persisted after the perturbation concluded.

Despite observing significant changes in the transient features of weight bearing (asymmetrical shifts in peak push-off vGRF and minimum midstance vGRF), we did not observe a change to vGRF impulse asymmetry, indicating that the relative contribution of each limb to body support during the full stance phase did not change. We can speculate that this may be because the perturbation we applied is mechanically efficient, despite the magnitude of disruption. For the walking frequencies we observed, the AdjuSST has a frequency response gain at or slightly above unity (25), meaning that nearly all of the mechanical energy put into the system by a participant’s weight is returned by the spring. For participants without neuromotor deficits, adaptation could involve modifying the transient aspects of weight-bearing while maintaining approximate symmetry in overall weight bearing impulse.

### Braking and propulsion aftereffects

Asymmetric stiffness walking elicited changes in peak braking force, but not in propulsive force. This finding runs counter to our hypothesis. As previously discussed, however, push-off vGRF and plantar flexor muscle activity shifted toward the perturbed side post-perturbation, suggesting that a motor adaptation increasing push-off overall was being elicited, but that the direction of the push-off force vector was aligned vertically. This contradicts a conclusion from (23) that increases in peak vGRF at pushoff imply increases in propulsion force, underscoring the need to directly measure braking and propulsion ground reaction forces when adaptation in propulsion is a relevant measure. This adaptation may result from the stiffness perturbation being constrained to the vertical direction. Likewise, the significant aftereffect in peak braking force asymmetry is coupled with a lack of significance in weight acceptance vGRF asymmetry, suggesting a more horizontal limb posture at heel-strike. It is possible that in order to achieve an increase in propulsion, the perturbation could be modified to additionally elicit an adaptation in trailing limb angle or by increasing propulsion demands overall (e.g., by introducing an upward incline) (29). It remains to be seen whether this is the case for walkers with chronic hemiparetic stroke.

Additionally, while we found a significant aftereffect in the peak braking force, it appears to rapidly wash out, nearly reaching symmetry within 10 strides (Figure 4). This rapid washout may be why we did not see a significant effect of condition in braking impulse, in which the first 5 strides and the last 5 strides of the post-early condition largely cancel out (Figure 4).

### Spatiotemporal symmetry was unaffected

Despite changes to ground reaction force and muscle excitation symmetry, we did not observe aftereffects in spatiotemporal symmetry. Using data from the force-sensing insoles, we measured a mean shift in stance time toward the unperturbed leg (∼1% asymmetry) and an increase in stride time by ∼30 ms during the perturbation, but we did not include these measurements in our statistical analysis because those were recorded at a lower temporal resolution and lower force sensing accuracy than the instrumented treadmill. Regardless of their statistical relevance, these changes are small compared to temporal changes observed during split-belt treadmill training (e.g., ∼30% at 4:1 belt speed ratio (30). This is possibly influenced by the belt speeds being tied in our perturbation, which may constrain the spatiotemporal gait parameters to remain closer to symmetry than if participants were allowed more freedom (i.e., on a self-paced treadmill or overground).

### Study limitations and future work

This study investigated the effects of asymmetric stiffness walking on healthy young adults to identify aftereffects indicative of neuromotor adaptation. While we were able to analyze our measures of interest before and after the perturbation, anterior-posterior GRF, step length, and stride length were not captured during the perturbation. We were able to measure EMG signals at full resolution during the perturbation, but vGRF were measured with force-sensing insoles which are not a perfect equivalent for the force plates in the rigid instrumented treadmill. Therefore, we included them to qualitatively illustrate the vGRF profile during the perturbation, but did not include them or the temporal measures computed from them in our analyses. Our protocol required transport of the participant between experimental setups before and after the perturbation. While we took precautions to prevent participants from taking any steps after exposure to the perturbation, the “Post-early” condition started 1-2 minutes after exposure. The first stride post-perturbation was also discarded to avoid including effects from the acceleration of the treadmill belts. Therefore, it is possible that the magnitude of the effects on gait immediately post-perturbation was not fully captured.

A limitation inherent to analysis of behavioral effects is that our conclusions about neural changes to motor control are indirect interpretations of our observations. This limitation can be addressed in part by replicating experimental results with neuromuscular models in which the mechanisms for motor control are fully described. In prior work, we used optimal control predictive simulations to describe split belt treadmill walking (31), and more recently to predict asymmetric stiffness walking to a wide range of mechanical parameters (32), in which many of the predicted changes to vGRF magnitude and muscle activity asymmetry were borne out in these experiments. Refinement of neuromuscular simulations focused on interpretation of our experimental findings will be an avenue for future work.

Finally, while insights into adaptation to asymmetric stiffness walking in healthy young adults are useful, they do not necessarily generalize to hemiparetic gait. As in the development of split belt treadmill training, this work quantifies the effect of the intervention in a healthy control population (17), a necessary step in the process of understanding the mechanisms that may eventually be leveraged toward clinical applications (16). We will investigate the effects of this intervention in individuals with chronic stroke in future.

## Conclusion

Asymmetric stiffness walking elicited changes to push-off GRF asymmetry corresponding with changes to plantar flexor activity, indicating a neuromotor adaptation to the perturbation. In contrast to split-belt treadmill perturbations, it did not elicit changes to spatiotemporal parameters of gait. Furthermore, while vertical GRF shifted toward the perturbed side limb during push-off, weight bearing impulse over the stance phase and propulsive force did not. Further study is needed to determine if asymmetric stiffness walking can elicit improved symmetry in push-off GRF in individuals with chronic stroke.

## Author Contributions

- Price: Conceived and designed research. Performed experiments. Analyzed data. Interpreted results of experiments. Prepared figures. Drafted manuscript. Edited and revised manuscript. Approved final version of manuscript.
- Schell: Performed experiments. Analyzed data. Interpreted results of experiments. Prepared figures. Drafted manuscript. Edited and revised the manuscript. Approved final version of the manuscript.
- Jaramillo: Performed experiments. Analyzed data. Interpreted results of experiments. Prepared figures. Approved final version of the manuscript.
- Chiasson: Performed experiments. Analyzed data. Interpreted results of experiments. Approved final version of the manuscript.
- Metsker: Performed experiments. Analyzed data. Interpreted results of experiments. Approved final version of the manuscript.
- Huber: Conceived and designed research. Edited and revised manuscript. Approved final version of manuscript.
- Hoogkamer: Conceived and designed research. Interpreted results of experiments. Drafted manuscript. Edited and revised manuscript. Approved final version of manuscript.

## Supplemental Figures

**Figure 1S.**
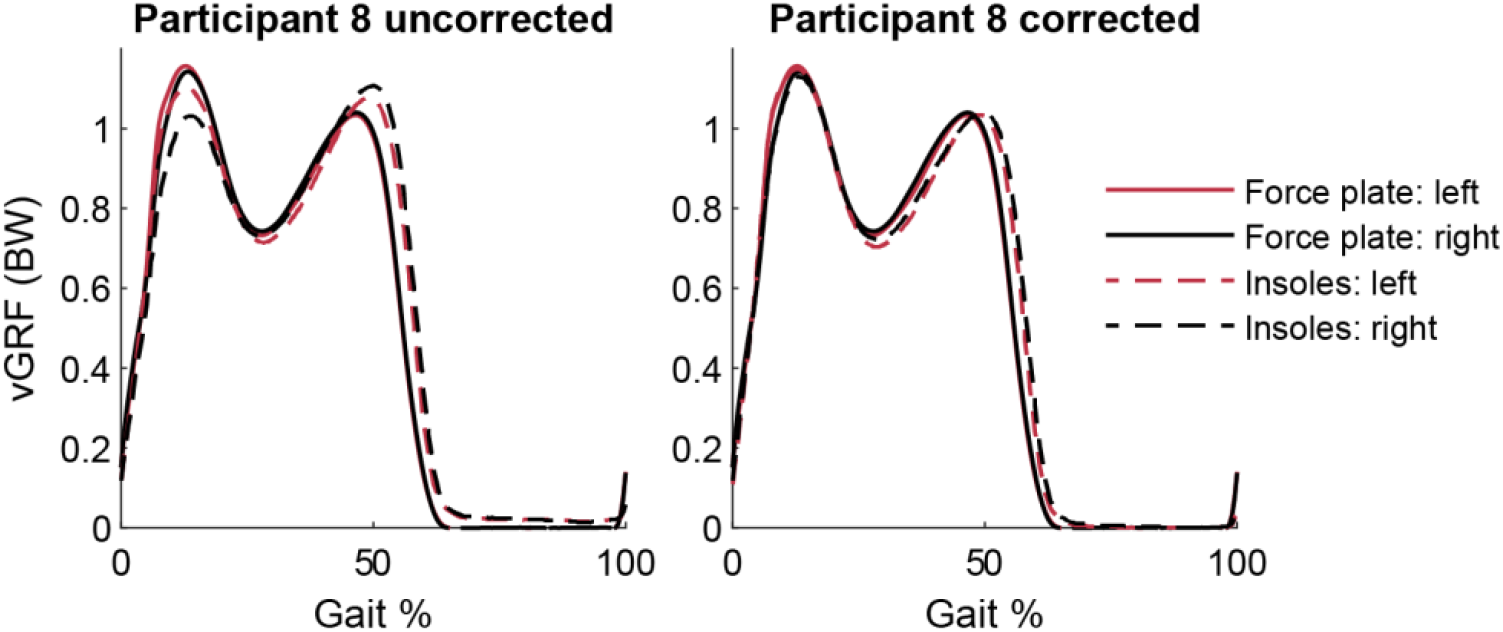
Uncorrected and corrected insole vertical GRF measurements compared with corresponding treadmill force plate vertical GRF measurements. Traces are the average of 20 strides during the baseline condition.

**Figure 2S.**
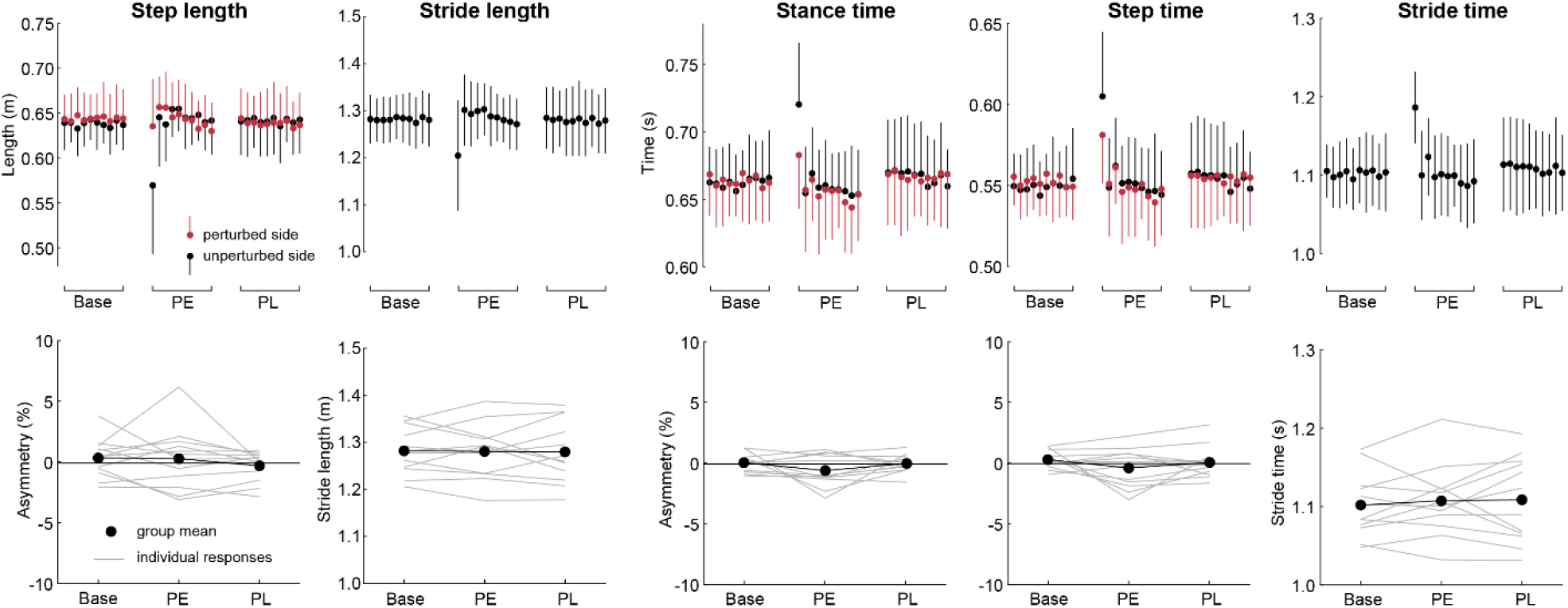
Group means of spatiotemporal measures per condition (Base = Baseline, PE = Post-early, PL = Post-late) (top) per stride and (bottom) asymmetry ratio, where relevant. Measures that comprise the sum of bilateral measures (stride length, stride time) are reported with their directly computed value rather than an asymmetry ratio. Error bars reflect one standard deviation. Asterisks reflect significant difference to baseline via paired t-test: ^*^ = p<.05, ^**^ = p<.01, ^***^ = p<.001.

## References

1. Alam Z, Rendos NK, Vargas AM, Makanjuola J, Kesar TM. Timing of propulsion-related biomechanical variables is impaired in individuals with post-stroke hemiparesis. Gait Posture 96: 275–278, 2022. doi: 10.1016/j.gaitpost.2022.05.022.

2. Hendrickson J, Patterson KK, Inness EL, McIlroy WE, Mansfield A. Relationship between asymmetry of quiet standing balance control and walking post-stroke. Gait Posture 39: 177–181, 2014. doi: 10.1016/j.gaitpost.2013.06.022.

3. Patterson KK, Parafianowicz I, Danells CJ, Closson V, Verrier MC, Staines WR, Black SE, McIlroy WE. Gait Asymmetry in Community-Ambulating Stroke Survivors. Arch Phys Med Rehabil 89: 304–310, 2008. doi: 10.1016/j.apmr.2007.08.142.

4. Patterson KK, Gage WH, Brooks D, Black SE, McIlroy WE. Evaluation of gait symmetry after stroke: A comparison of current methods and recommendations for standardization. Gait Posture 31: 241–246, 2010. doi: 10.1016/j.gaitpost.2009.10.014.

5. Ugur C, Gücüyener D, Uzuner N, Ozkan S, Ozdemir G. Characteristics of falling in patients with stroke. J Neurol Neurosurg Psychiatry 69: 649–651, 2000. doi: 10.1136/jnnp.69.5.649.

6. Mijic M, Schoser B, Young P. Efficacy of functional electrical stimulation in rehabilitating patients with foot drop symptoms after stroke and its correlation with somatosensory evoked potentials—a crossover randomised controlled trial. Neurol Sci 44: 1301–1310, 2023. doi: 10.1007/s10072-022-06561-3.

7. Nadeau S, Gravel D, Arsenault AB, Bourbonnais D. Plantarflexor weakness as a limiting factor of gait speed in stroke subjects and the compensating role of hip flexors. Clin Biomech 14: 125–135, 1999. doi: 10.1016/s0268-0033(98)00062-x

8. Roelker SA, Bowden MG, Kautz SA, Neptune RR. Paretic propulsion as a measure of walking performance and functional motor recovery post-stroke: A review. Gait Posture 68: 6–14, 2019. doi: 10.1016/j.gaitpost.2018.10.027

9. Combs-Miller SA, Kalpathi Parameswaran A, Colburn D, Ertel T, Harmeyer A, Tucker L, Schmid AA. Body weight-supported treadmill training vs. overground walking training for persons with chronic stroke: A pilot randomized controlled trial. Clin Rehabil 28: 873–884, 2014. doi: 10.1177/0269215514520773.

10. Turnbull GI, Charteris J, Wall JC. A comparison of the range of walking speeds between normal and hemiplegic subjects. Scand J Rehabil Med 3: 175–182, 1995. doi: 10.2340/165019771995175182

11. Titianova EB, Pitkänen K, Pääkkönen A, Sivenius J, Tarkka IM. Gait characteristics and functional ambulation profile in patients with chronic unilateral stroke. Am J Phys Med Rehabil 82: 778–786, 2003. doi: 10.1097/01.PHM.0000087490.74582.E0

12. Bonnyaud C, Pradon D, Zory R, Bensmail D, Vuillerme N, Roche N. Does a single gait training session performed either overground or on a treadmill induce specific short-term effects on gait parameters in patients with hemiparesis? a randomized controlled study. Top Stroke Rehabil 20: 509–518, 2013. doi: 10.1310/tsr2006-509.

13. Sackley C. The relationships between weight-bearing asymmetry after stroke, motor function and activities of daily living. Physiother Theory Pract 6: 179–185, 1990. doi: 10.3109/09593989009048293

14. Li W, Li Y, Gao Q, Liu J, Wen Q, Jia S, Tang F, Mo L, Zhang Y, Zhai M, Chen Y, Guo Y, Gong W. Change in knee cartilage components in stroke patients with genu recurvatum analysed by zero TE MR imaging. Sci Rep 12, 3751, 2022. doi: 10.1038/s41598-022-07817-w.

15. Jørgensen L, Jacobsen BK, Wilsgaard T, Magnus JH. Walking after Stroke: Does It Matter? Changes in Bone Mineral Density Within the First 12 Months after Stroke. A Longitudinal Study. Osteoporos Int 11: 381–387, 2000.

16. Reisman DS, McLean H, Keller J, Danks KA, Bastian AJ. Repeated split-belt treadmill training improves poststroke step length asymmetry. Neurorehabil Neural Repair 27: 460–468, 2013. doi: 10.1177/1545968312474118.

17. Reisman DS, Block HJ, Bastian AJ. Interlimb coordination during locomotion: What can be adapted and stored? J Neurophysiol 94: 2403–2415, 2005. doi: 10.1152/jn.00089.2005.

18. Awad LN, Lewek MD, Kesar TM, Franz JR, Bowden MG. These legs were made for propulsion: advancing the diagnosis and treatment of post-stroke propulsion deficits. J Neuroeng Rehabil 17: 139, 2020. doi: 10.1186/s12984-020-00747-6

19. Lewek MD, Braun CH, Wutzke C, Giuliani C. The role of movement errors in modifying spatiotemporal gait asymmetry post stroke: a randomized controlled trial. Clin Rehabil 32: 161–172, 2018. doi: 10.1177/0269215517723056.

20. Santucci V, Alam Z, Liu J, Spencer J, Faust A, Cobb A, Konantz J, Eicholtz S, Wolf S, Kesar TM. Immediate improvements in post-stroke gait biomechanics are induced with both real-time limb position and propulsive force biofeedback. J Neuroeng Rehabil 20, 2023. doi: 10.1186/s12984-023-01154-3.

21. Hinton EH, Buffum R, Kingston D, Stergiou N, Kesar T, Bierner S, Knarr BA. Real-Time Visual Kinematic Feedback During Overground Walking Improves Gait Biomechanics in Individuals Post-Stroke. Ann Biomed Eng 52: 355–363, 2024. doi: 10.1007/s10439-023-03381-0.

22. Kal E, Winters M, van der Kamp J, Houdijk H, Groet E, van Bennekom C, Scherder E. Is Implicit Motor Learning Preserved after Stroke? A Systematic Review with Meta-Analysis. PLoS One 11: e0166376, 2016. doi: 10.1371/journal.pone.0166376.

23. Chambers V, Artemiadis P. Using robot-assisted stiffness perturbations to evoke aftereffects useful to post-stroke gait rehabilitation. Front Robot AI 9, 2023. doi: 10.3389/frobt.2022.1073746.

24. Winter DA. The biomechanics and motor control of human gait. University of Waterloo Press, 1987. doi: 10.1002/9780470549148

25. Price M, Locurto D, Abdikadirova B, Huber ME, Hoogkamer W. Design and Characterization of the AdjuSST: An Adjustable Surface Stiffness Treadmill. IEEE/ASME Transactions on Mechatronics, 2024 (Early access). doi: 10.1109/TMECH.2024.3508476

26. Shiavi R, Frigo IC, Pedotti A. Electromyographic signals during gait: criteria for envelope filtering and number of strides. Medical and Biological Engineering and Computing 36: 171–178, 1998. doi: 10.1007/BF02510739

27. Emanuel Singh R, Iqbal K, White G, K. Holtz J. A Review of EMG Techniques for Detection of Gait Disorders. In: Artificial Intelligence - Applications in Medicine and Biology, edited by Aceves-Fernandez MA. IntechOpen, 2019. doi: 10.5772/intechopen.84403

28. Lamontagne A, Malouin F, Richards CL, Dumas F. Mechanisms of disturbed motor control in ankle weakness during gait after stroke. Gait Posture 15: 244–255, 2002. doi: 10.1016/S0966-6362(01)00190-4.

29. Sombric CJ, Torres-Oviedo G. Augmenting propulsion demands during split-belt walking increases locomotor adaptation of asymmetric step lengths. J Neuroeng Rehabil 17: 69, 2020. doi: 10.1186/s12984-020-00698-y.

30. Stenum J, Choi JT. Step time asymmetry but not step length asymmetry is adapted to optimize energy cost of split-belt treadmill walking. J Physiol 598: 4063–4078, 2020. doi: 10.1113/JP279195.

31. Price M, Huber ME, Hoogkamer W. Minimum effort simulations of split-belt treadmill walking exploit asymmetry to reduce metabolic energy expenditure. J Neurophysiol 129: 900–913, 2023. doi: 10.1152/jn.00343.2022.

32. Abdikadirova B*, Price M*, Hoogkamer W, Huber ME. Forward Simulations of Walking on Surfaces with Asymmetric Mechanical Impedance: Insights for Gait Rehabilitation. Proc Int Conf IEEE Eng Med Bio Soc (accepted), 2025. Preprint doi: 10.1101/2024.10.03.616487

